# Bacterial Sphingolipids Exacerbate Colitis by Inhibiting ILC3-derived IL-22 Production

**DOI:** 10.1101/2023.09.05.555400

**Authors:** Bin Bao, Youyuan Wang, Pavl Boudreau, Xinyang Song, Meng Wu, Xi Chen, Izabel Patik, Ying Tang, Jodie Ouahed, Amit Ringel, Jared Barends, Chuan Wu, Emily Balskus, Jay Thiagarajah, Jian Liu, Michael R. Wessels, Wayne Lencer, Dennis L. Kasper, Dingding An, Bruce Horwitz, Scott B. Snapper

**Author notes:** Correspondence: Bin Bao or Scott B. Snapper Boston Children’s Hospital, and Harvard Medical School, Boston, MA, 02115.

## Abstract

Commensal bacteria of the Bacteroidetes phylum are the primary producers of sphingolipids in the gut lumen. These lipids serve dual roles as bacterial virulence factors and regulators of the host mucosal immune system, including regulatory T cells and invariant natural killer T cells (iNKT). Sphingolipid composition is significantly altered in fecal samples of patients with inflammatory bowel disease (IBD). However, the specific mechanisms by which bacterial sphingolipids modulate mucosal homeostasis and regulate intestinal inflammation remain unclear. In this study, we investigated the impact of bacterial sphingolipids on intestinal inflammation by mono-colonizing mice with *Bacteroides fragilis* strains that either express or lack sphingolipids during DSS-induced colitis. We discovered that *B. fragilis* sphingolipids exacerbate intestinal inflammation. Mice mono-colonized with *B. fragilis* lacking sphingolipids exhibited less severe DSS-induced colitis. This amelioration of colitis was associated with increased production of interleukin-22 (IL-22) by innate lymphoid cell type 3 (ILC3). Consistent with the inhibitory effect of sphingolipids on IL-22 production, mice colonized with *B. fragilis* lacking sphingolipids showed enhanced epithelial STAT3 activity, intestinal cell proliferation, and antimicrobial peptide production following DSS treatment compared to those colonized with *B. fragilis* producing sphingolipids. Additionally, colitis severity in mice colonized with *B. fragilis* lacking sphingolipids was exacerbated upon IL-22 blockade. Furthermore, our study reveals that bacterial sphingolipids restrict epithelial IL-18 production following DSS treatment and interfere with IL-22 production by a subset of ILC3 cells expressing both the interleukin-18 receptor (IL-18R) and major histocompatibility complex class II (MHC II). These findings indicate that *B. fragilis*-derived sphingolipids exacerbate mucosal inflammation by impeding epithelial IL-18 expression, resulting in compromised production of IL-22 by ILC3 cells.

**Highlights:**

1. *B. fragilis*-derived sphingolipids exacerbate DSS-induced colitis in mono-colonized C57BL/6 mice.
2. *B. fragilis*-derived sphingolipids constrain ILC3-derived IL-22, leading to reduced colonic epithelial cell proliferation and compromised barrier function.
3. *B. fragilis*-derived sphingolipids restrict epithelial NLRC4 inflammasome activation and IL-18 secretion.
4. *B. fragilis*-derived sphingolipids modulate IL-22 production by IL18R^+^ MHC II^+^ ILC3s.

## Introduction

The commensal microbiota plays a critical role in maintaining intestinal homeostasis and overall health. Microbial metabolites have been recognized for their diverse functions in regulating mucosal homeostasis^1–3^. Inflammatory bowel disease (IBD) patients exhibit alterations in gut microbial metabolites^4–6^, highlighting the potential importance of understanding the specific roles of microbial products in IBD pathogenesis^6,7^.

Sphingolipids, essential bioactive and structural molecules, have emerged as key regulators of metabolic disorders and immunological responses in mammals^8,9^. Sphingolipids levels are tightly controlled through various mechanisms, including *de novo* synthesis, recycling, and intestinal uptake. In addition to dietary sources, sphingolipids are introduced into the intestinal lumen by certain commensal bacteria. Intestinal bacteria belonging to the Bacteroidetes phylum produce sphingolipids through the enzyme serine palmitoyltransferase (SPT)^10^. Among the human intestinal microbiota, Bacteroidetes are the primary producers of sphingolipids^11,12^. Microbial metabolites of *Bacteroides fragilis*, one of the most abundant commensal microbes, have been extensively studied for their interaction with the host immune^13,14^.

*B. fragilis*-derived sphingolipids interact with host epithelial and immune cells, influencing host lipid metabolism^15^ and modulating mucosal immune cells, including iNKT cells^13,16,17^. One specific sphingolipid, *B. fragilis* α-Galactosyl ceramide (Bfa-GL-Cers), exhibits structural similarity to KRN7000, the prototypical agonist of iNKT cells^12^. Interestingly, Bfa-GL-Cers has demonstrated both stimulatory^13^ and regulatory^16^ effects on iNKT cells. Bfa-GL-Cers also suppresses intestinal serotonin secretion and associated gut motility^18^. The presence of structural variations in branched sphinganine chains may explain the paradoxical immunomodulatory effects of Bfa-GL-Cers on iNKT cells^17^. Sphingolipid-dependent regulation of iNKT cells modulates the susceptibility of oxazolone-induced colitis^12,16,19^. Moreover, sphingolipids have been identified as the most abundant fecal metabolite class in patients with IBD^20^. In the context of IBD, among various microbial sphingolipids, *B. fragilis*-derived sphingolipids are particularly intriguing due to their interactions with host immune cells, including iNKT cells. However, the functional role of these sphingolipids and their roles in IBD and other mucosal immune cells beyond iNKT cells remain incompletely understood.

Innate lymphoid cells (ILCs) are essential components of the innate immune system, residing in tissues and playing a critical role in regulating host-microbe interactions and maintaining mucosal barrier function^21–23^. Among the ILC subsets, group 3 ILCs (ILC3s) are highly enriched in mucosal tissues and have been implicated in cellular interactions within the intestine, both in healthy and diseased states^22,24,25^. The production of IL-22 by ILC3s is vital for promoting intestinal epithelial cell proliferation and the synthesis of antimicrobial peptides, which is crucial for maintaining the integrity of the intestinal barrier^26–28^. Furthermore, ILC3s have the ability to interact with CD4^+^ T cells through major histocompatibility complex class II (MHC II) molecules, suppressing commensal bacteria-specific CD4^+^ T cell responses^29^. Consequently, ILC3s mediate communication between the microbiota and the host mucosal immune system. Importantly, studies have demonstrated that IL-22 produced by ILC3s is protective in reducing the severity of dextran-sodium sulfate (DSS) colitis. IL-22 has beneficial effects by activating STAT3 in intestinal epithelial cells, enhancing proliferation and barrier function, thus promoting mucosal healing^30^. Despite the observation of reduced ILC3 numbers or subsets and impaired effector functions in inflamed tissues of individuals with IBD^29,31,32^, the specific mechanisms underlying the response of ILC3s to commensal microbial metabolites, including sphingolipids, remain poorly understood. Unraveling the intricate interactions between ILC3s and microbial products, such as *B. fragilis*-derived sphingolipids, may provide critical insights into the pathogenesis of IBD and potential therapeutic strategies for targeting this essential immune cell population.

In this study, we have identified a role for *B. fragilis*-derived sphingolipids in modulating mucosal immune responses through ILC3s. We found that *B. fragilis*-derived sphingolipids impair epithelial cell proliferation and exacerbate DSS-induced colitis in germ-free mice mono-colonized with *B. fragilis* strains. *B. fragilis*-derived sphingolipids inhibit the IL-22-STAT3 axis and regulate the NLRC4 inflammasome and IL-18 secretion in epithelial cells. Additionally, a specific subset of ILC3s expressing IL-18R and MHC II responds to these sphingolipids, modulating IL-22 expression. These findings highlight the intricate interplay between *B. fragilis*-derived sphingolipids, host epithelial cells, and ILC3s that modulate intestinal inflammation and homeostasis.

## Results

### *B. fragilis*-derived sphingolipids exacerbate DSS-induced colitis

To explore the function of *B. fragilis*-derived sphingolipids in the gut, we employed a model in which mice were mono-colonized with either wild-type *B. fragilis* or a mutant strain that lacks serine palmitoyltransferase (SPT), the enzyme that catalyzes the first step in sphingolipid biosynthesis, resulting in the absence of sphingolipids^16^. Germ-free C57BL/6 mice were mono-colonized with either wild-type or mutant *B. fragilis*, generating two distinct strains: BFWT mice or BFΔSPT mice. Both strains of mono-colonized mice exhibited comparable growth rates (Figure S1A) and fecal colony forming units (CFUs) at twelve weeks of age (Figure S1B), indicating that bacterial sphingolipids do not significantly alter weight gain or *B. fragilis* colonization in this mono-colonized setting. In addition, neither BFWT nor BFΔSPT adult mice showed signs of intestinal or systemic inflammation, as evidenced by similar colon lengths and comparable weights of mesenteric lymph nodes (mLN) and spleen (Figure S1C-E).

To investigate the role of *B. fragilis*-derived sphingolipids in the development of colitis, mono-colonized mice (BFWT and BFΔSPT) were exposed to dextran sodium sulfate (DSS), a substance that induces acute epithelial injury and colitis. BFWT mice exposed to DSS exhibited more severe colitis symptoms compared to BFΔSPT mice. BFWT mice showed increased weight loss, diarrhea, rectal bleeding, and splenic weight (Figure 1A-C, Figure S1G), along with shorter colons and higher histopathology inflammation scores (Figure 1D-G), indicating more severe inflammation.

**Figure 1:**
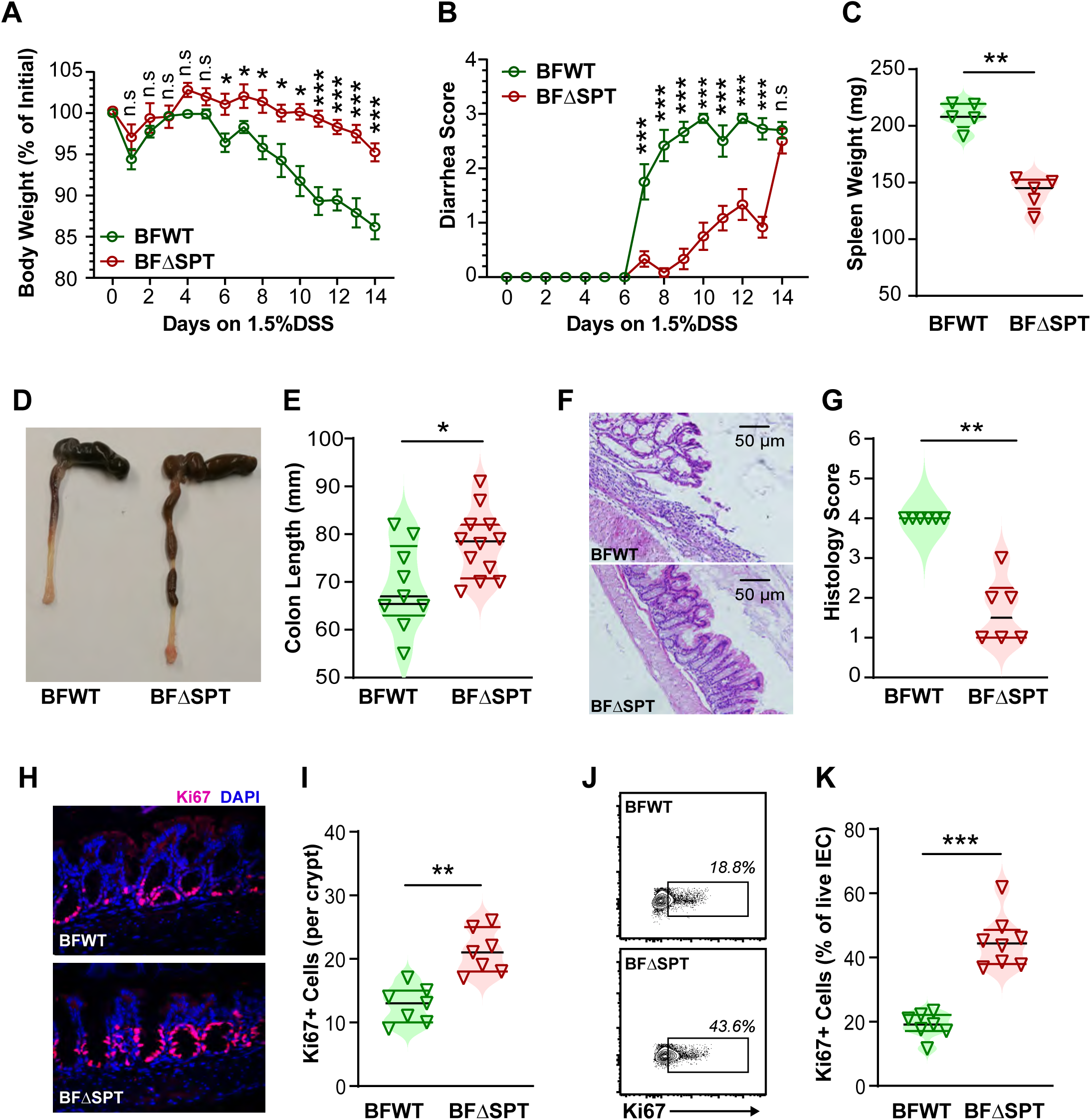
*B. fragilis*-derived sphingolipids exacerbate DSS-induced colitis. (**A**) Weight loss and (**B**) diarrhea scores were measured in mono-colonized mice treated with 1.5% DSS in their drinking water for 14 days. (**C**) Spleen weight and (**D-E**) colons length were measured on day 14. (**F**) Histopathology scores of the colon were assessed, and representative (**G**) H&E-stained colon sections were examined. (**H**) Representative immunofluorescence sections of the colon showing Ki67 (pink) and DAPI (blue) staining were analyzed. (**I**) Quantification of Ki67^+^ cells number per crypt. (**J**) Representative flow cytometry plots and (**K**) quantification of colonic Ki67^+^ epithelial cells based on the flow cytometry results. IECs were gated on singlet/live/CD45-/EpCAM^+^ cells. The data represent two or three independent experiments and are shown as means ± s.e.m. Statistical significance was determined using the nonparametric Mann-Whitney test: *p<0.05, **p<0.01, ***p<0.001.

To explore how *B. fragilis*-derived sphingolipids exacerbate colitis, we examined whether they affect colonic epithelial proliferation using Ki67 immunofluorescence staining and flow cytometric analyses. Upon exposure to DSS, BFWT mice exhibited fewer proliferating colonic epithelial cells than BFΔSPT mice, as indicated by fewer Ki67^+^ cells (Figure 1H-K). These results suggest that *B. fragilis*-derived sphingolipids may exacerbate DSS-induced colitis potentially by restricting the proliferative capacity of colonic epithelial cells following (or during) injury.

### *B. fragilis*-derived sphingolipids restrain colonic STAT3 activation in colonic epithelial cells

To further investigate the inhibitory effect of *B. fragilis*-derived sphingolipids on colonic epithelial cell proliferation, we conducted experiments using murine colonic organoids. These organoids were exposed to total lipids extracted from either WT or SPT mutant *B. fragilis* strains, or purified sphingolipids obtained from *B. fragilis*, to directly assess their impact on proliferation. Neither total lipids nor purified sphingolipids from *B. fragilis* demonstrated any significant influence on organoids growth, as evidenced by the comparable number of spheres and average sphere area in each Matrigel dome (Figure S2A-C). Consistent with these organoid growth results, the percentage of Ki67+ cells in organoids remained unaltered upon treatment with *B. fragilis*-derived sphingolipids (Figure S2D-E). Thus, *B. fragilis*-derived sphingolipids do not directly regulate colonic epithelial cell proliferation, suggesting the involvement of other cells or cytokines in the sphingolipid-dependent regulation of colonic epithelial cell proliferation.

To gain further insight into the mechanism underlying the effect of *B. fragilis* sphingolipids on epithelial cell function after DSS challenge, we performed RNA sequencing analysis of colonic epithelial cells from DSS-treated BFWT and BFΔSPT mice. We identified 508 differentially expressed genes (padj < 0.05), with 38 genes up-regulated and 52 genes down-regulated by more than twofold in BFΔSPT mice compared to BFWT mice (Supplementary Table 1-3). Among these genes, *Reg3b*/*Reg3g*, and *Mmp13* were significantly increased in BFΔSPT mice (Figure 2A). Reg3b/Reg3g encode proteins associated with epithelial barrier function^33,34^, and Mmp13 regulates wound healing^35^. These results were further validated by quantitative PCR (Figure S3A). *Reg3b*/*Reg3g*, and *Mmp13* are known to be regulated by signal transducer and activator of transcription 3 (STAT3), whose activity is crucial for mucosal wound healing following DSS challenge^30^.

**Figure 2:**
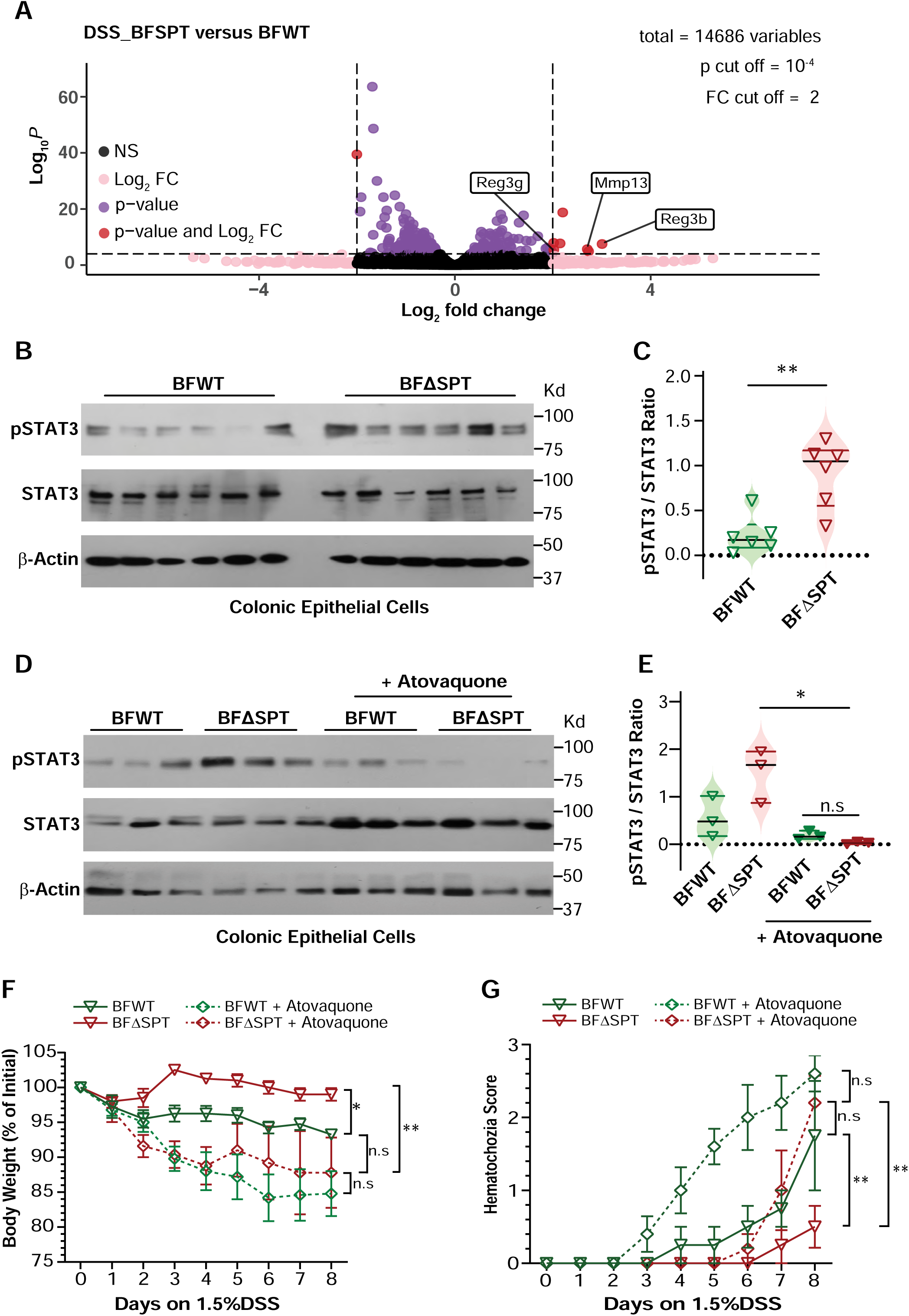
*B. fragilis*-derived sphingolipids suppress STAT3 in colonic epithelial cells. (**A**) Volcano plot showing differentially expressed genes in colonic epithelial cells from mono-colonized mice treated with 1.5% DSS in drinking water for 14 days. (**B**) Immunoblot analysis and (**C**) corresponding quantification of phosphorylated and total STAT3 in colonic epithelial cells from mono-colonized mice treated with 1.5% DSS in drinking water for 14 days (n = 6 mice per group). (**D**) Immunoblot analysis and (**E**) corresponding quantification of phosphorylated and total STAT3 in colonic epithelial cells from mono-colonized mice treated with 1.5% DSS in drinking water for 14 days (n = 3 mice per group). Two groups of mice were orally gavaged with atovaquone (a STAT3 inhibitor) or PBS every other day. (**F**) Weight loss and (**G**) Hematochezia scores were monitored during the 8-day DSS treatment in mono-colonized mice that received atovaquone or PBS gavage. The data represent means ± s.e.m. of two or three independent experiments. Statistical significance was determined using the nonparametric Mann-Whitney test: *p<0.05, **p<0.01, ***p<0.001.

To investigate the effect of *B. fragilis*-derived sphingolipids on STAT3 activity, we compared the levels of phosphorylated STAT3 (pSTAT3) in colonic epithelial cells isolated from DSS-treated BFWT and BFΔSPT mice by Western Blot analysis. Colonic epithelial cells from BFΔSPT mice exhibited significantly higher pSTAT3 activity compared to BFWT mice (Figure 2B-C). To further explore whether enhanced STAT3 activation observed in BFΔSPT mice confers protection against DSS-induced colitis, we treated monocolonized mice with the STAT3 inhibitor atovaquone. As expected, atovaquone suppressed STAT3 activity in the colonic epithelial cells of both DSS-treated BFWT and BFΔSPT mice (Figure 2D-E). The inhibition of STAT3 resulted in more severe colitis in both DSS-treated BFWT and BFΔSPT mice, and with STAT3 inhibition significant differences in several measures of intestinal inflammation, including body weight loss (Figure 2F), diarrhea (Figure S3B), and rectal bleeding scores (Figure 2G) were no longer observed between these groups. These results suggest that *B. fragilis*-derived sphingolipids modulate intestinal inflammation, by inhibiting colonic epithelial STAT3 activation.

### *B. fragilis*-derived sphingolipids modulate the colonic IL-22-STAT3 axis to augment DSS-induced colitis

To investigate whether *B. fragilis*-derived sphingolipids directly modulate epithelial STAT3 activation, murine colonic organoids were treated with sphingolipids purified from *B. fragilis*. Intriguingly, the activation of phosphorylated STAT3 was not affected following short-term or long-term treatment with either total lipids or purified sphingolipids from *B. fragilis* (Figure S4). These data suggest that modulation of colonic STAT3 activity by *B. fragilis* sphingolipids is an indirect effect. To explore this further, cytokines known to mediate epithelial STAT3 activity, specifically IL-6, IL-10, and IL-22, were examined in colonic explants from DSS-treated BFWT and BFΔSPT mice. Remarkably, BFΔSPT mice had higher levels of colonic IL-22 than BFWT mice, with no significant difference in IL-6 and IL-10 levels (Figure 3A-C).

**Figure 3:**
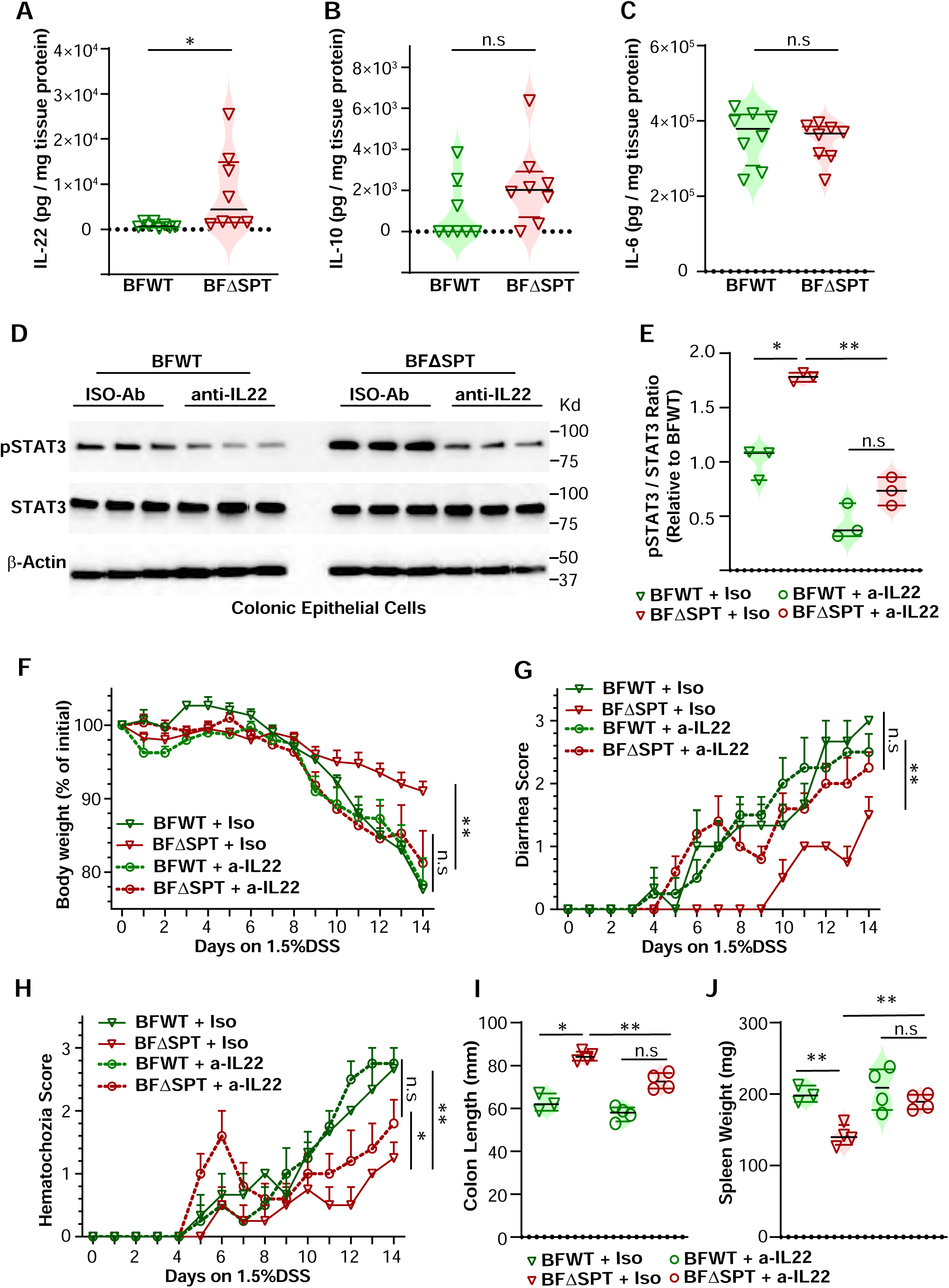
*B. fragilis*-derived sphingolipids suppress STAT3 by constraining the IL-22 signal. (**A-C**) Concentrations of IL-22, IL-10, and IL-6 were measured by ELISA in the medium of colon explant cultures from mono-colonized mice challenged with 1.5% DSS in drinking water for 14 days. Protein extraction was performed from the colon explants after culture, and cytokine concentrations were normalized to protein weight (n = 8 individual mice per group). (**D-J**) Mono-colonized mice were treated with 1.5% DSS in drinking water for 14 days and injected with either anti-IL-22 or Isotype IgG. (**D**) Immunoblot analysis and (**E**) quantification of phosphorylated and total STAT3 in colonic epithelial cells from mono-colonized mice treated with 1.5% DSS in drinking water for 14 days (n = 3 mice per group). (**F**) Weight loss, (**G**) diarrhea scores and (**H**) hematochezia scores were recorded during the DSS treatment. (**I**) Colon length and (**J**) spleen weight was assessed on day 14 (n = 3-4 individual mice per group). The data represent means ± s.e.m. of two or three independent experiments. Statistical significance was determined using the nonparametric Mann-Whitney test: *p<0.05, **p<0.01, ***p<0.001.

To further explore the role of IL-22 in modulating host epithelial STAT3 activity and colitis pathogenesis in response to *B. fragilis*-derived sphingolipids, we administered IL-22 antibodies to monocolonized mice challenged with DSS. As expected, IL-22 blockade significantly reduced STAT3 activity in the colonic epithelium of both BFΔSPT and BFWT mice (Figure 3D-E). Further, IL-22 blockade caused increased colitis severity in DSS-treated BFΔSPT mice, resulting in comparable severity of colitis in BFΔSPT and BFWT mice (Figure 3F-J). Taken together, our results suggest that *B*. *fragilis*-derived sphingolipids restrain the colonic IL-22 production and epithelial STAT3 activity, thereby exacerbating DSS-induced colitis.

### *B. fragilis* sphingolipids constrain ILC3s-derived IL-22 and epithelial cell-derived IL-18

To identify the specific IL-22-producing cells affected by *B. fragilis*-derived sphingolipids, we analyzed lamina propria cells isolated from DSS-treated BFWT and BFΔSPT mono-colonized mice. Colonic ILC3s from DSS-treated BFΔSPT mice exhibited enhanced IL-22 secretion compared with those from BFWT mice (Figure 4A-B). In contrast, no significant difference in IL-22 expression was observed among other cell types implicated in intestinal IL-22 production, including gamma-delta T (γδ T) cells, natural killer (NK) cells, and T Helper 17 (Th17) cells (Figure S5). These results suggest that *B. fragilis*-derived sphingolipids modulate IL-22 production in colonic ILC3s.

**Figure 4:**
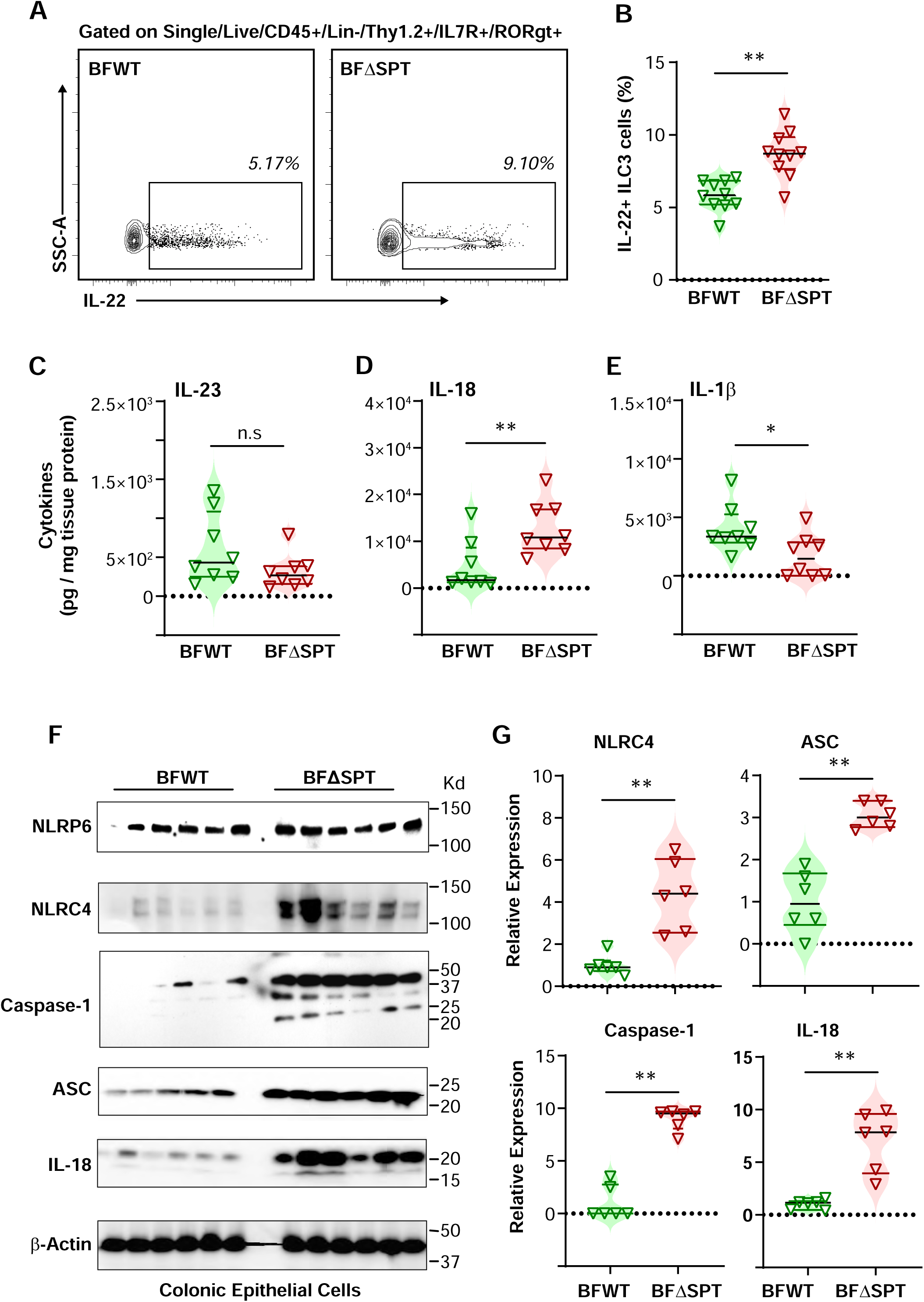
*B. fragilis*-derived sphingolipids suppress colonic IL-22 and IL-18 activity by interacting with ILC3s and epithelial cells. (**A**) Representative flow cytometry plots and (**B**) quantification of IL-22^+^ ILC3 cells in the colon of mono-colonized mice challenged with 1.5% DSS in drinking water for 7 days. (**C-E**) Concentrations of IL-23, IL-18, and IL-1β were measured in the medium of colon explant cultures from mono-colonized mice challenged with 1.5% DSS in drinking water for 7 days. Protein extraction was performed from the colon explants after culture, and cytokine concentrations were normalized to protein weight (n = 8 individual mice per group). (**F**) Immunoblot analysis and (**G**) quantification of IL-18 and inflammasome proteins in colonic epithelial cells from mono-colonized mice treated with 1.5% DSS in drinking water for 7 days (n = 6 mice per group). Data represent two or three independent experiments shown as the means± s.e.m. Statistical significance was determined by the nonparametric Mann-Whitney test. *p < 0.05, **p < 0.01, ***p < 0.001.

The production of IL-22 in ILC3 cells is regulated by several cytokines, including IL-23, IL-1β, and IL-18, which are known to induce IL-22 production in ILC3 cells during DSS-induced colitis^36,37^. To determine whether *B. fragilis*-derived sphingolipids inhibit the expression of any of these cytokines, we further investigated the key intestinal cytokines associated with the regulation of IL22 production by ILC3 cells in DSS-treated BFWT and BFΔSPT mono-colonized mice. Interestingly, BFΔSPT mice had higher levels of colonic IL-18 when compared to BFWT mice (Figure 4C-D), with no significant increase in either IL23 or IL-1β in BFΔSPT mice (Figure 4E). Thus, IL-23 and IL-1β are not responsible for the elevated ILC3-derived IL-22 when *B. fragilis* sphingolipids are absent in the mouse intestine. These results suggest that *B. fragilis*-derived sphingolipids inhibit the ability of ILC3 to produce IL-22 by inhibiting the production of IL-18.

During acute DSS-induced colitis, inflammasome activation in intestinal epithelial cells and subsequent IL-18 secretion drive IEC proliferation and tissue regeneration, limiting mucosal damage and immune cell activation in the lamina propria^38,39^. Therefore, we asked whether the level of inflammasome components, including Caspase-1, Nlrp6, and Nlrc4, differed within the colonic epithelium of BFWT and BFΔSPT mono-colonized mice challenged with DSS. Consistent with the ELISA results, IL-18 levels measured by western blot were higher in the colonic epithelial cells of BFΔSPT mice challenged with DSS than in BFWT mice (Figure 4F-G). Moreover, NLRC4 and Caspase 1 were significantly increased in colonic epithelial cells of BFΔSPT mice challenged with DSS compared to BFWT mice (Figure 4F-G). However, colonic epithelial NLRP6 protein levels were comparable between the two DSS-treated mono-colonized mice (Figure 4F-G). In addition, NLRP3 was not detectable in the epithelial cells of the mono-colonized mice. These results suggest that *B. fragilis*-derived sphingolipids suppress NLRC4 and Caspase-1, leading to impaired IL-18 maturation in colonic epithelial cells during DSS-induced colitis.

### *B. fragilis-*derived sphingolipids modulate ILC3-derived IL-22 by targeting IL18R^+^ MHCII^+^ ILC3

Given the altered IL18 expression and ILC3-derived IL22 expression in response to *B. fragilis*-derived sphingolipids, we sought to evaluate the expression of the IL-18 receptor (IL-18R) on colonic ILC3s. Importantly, as previously described^40^, IL-22 is predominantly expressed in colonic IL-18R^+^ ILC3s (Figure S6A). The percentage of colonic IL-18R^+^ ILC3s was comparable between the BFWT and BFΔSPT mice challenged with DSS (Figure 5A-B). Although not statistically significantly different (p = 0.0635), there was a trend of enhanced IL-22 expression in the IL-18R^+^ ILC3s of BFΔSPT mice challenged with DSS (Figure 5C-D).

**Figure 5:**
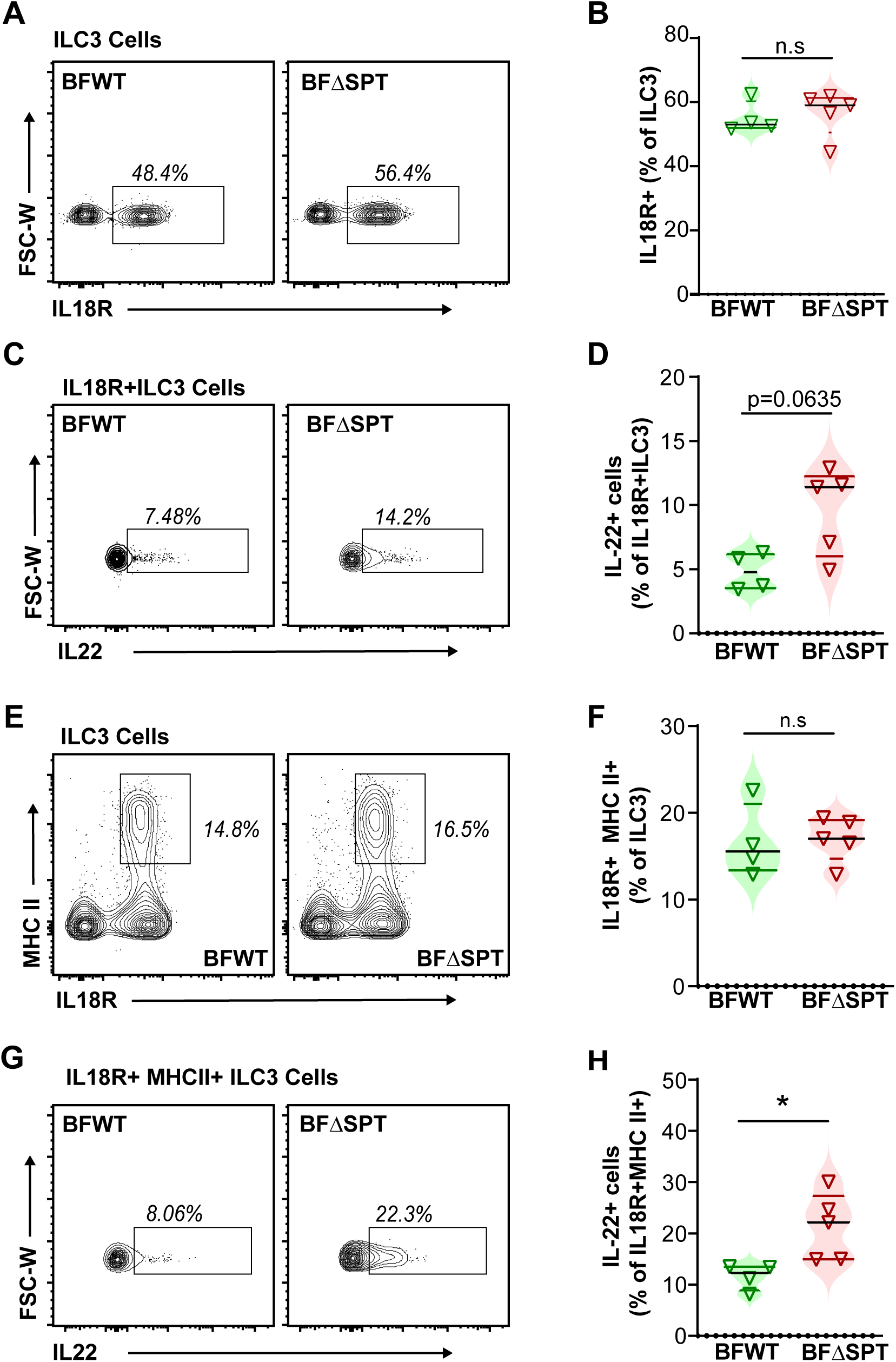
*B. fragilis*-derived sphingolipids modulate ILC3-derived IL-22 by targeting IL18R+ MHCII+ ILC3 cells. (**A**) Representative flow cytometry plots and (**B**) quantification of IL-18R^+^ cells in ILC3 cells in the colon of mono-colonized mice after 7 days of 1.5% DSS challenge. (**C**) Representative flow cytometry plots and (**D**) quantification of IL-22^+^ cells in IL-18R^+^ ILC3 cells in the colon of mono-colonized mice after 7 days of 1.5% DSS challenge. (**E**) Representative flow cytometry plots and quantification of (**F**) IL18R+ MHCII^+^ cells in ILC3 cells in the colon of mono-colonized mice after 7 days of 1.5% DSS challenge. (**G**) Representative flow cytometry plots and quantification of (**H**) IL-22^+^ cells in IL18R^+^MHCII^+^ ILC3 cells in the colon of mono-colonized mice after 7 days of 1.5% DSS challenge. ILC3 cells are gated on single/live/CD45^+^/Lin-/Thy1.2^+^/IL7R^+^/RORγt^+^ cells. A-C, n = 8 individual mice per group; D-M, n = 4-5 individual mice per group. Data represents means± s.e.m. from two or three independent experiments. The nonparametric Mann-Whitney test determined statistical significance (*p<0.05, **p<0.01, ***p<0.001).

To further investigate the regulation of IL-22 production in ILC3s, we examined the expression of MHC-II on colonic ILC3s, which has been reported to play a role in regulating IL-22 production in ILC3s^21,24^. MHC II^+^ ILC3 cells produced more IL-22 than MHC II^-^ ILC3s in mono-colonized mice (Figure S6B). However, the percentage of colonic MHC II^+^ ILC3s, as well as the fraction of these cells that expressed IL-22, was comparable between BFWT and BFΔSPT mice challenged with DSS (Figure S6C-F).

While the percentage of colonic IL-18R^+^ MHC II^+^ ILC3s was equivalent between BFWT and BFΔSPT mice (Figure 5E-F), the percentage of these cells that expressed IL-22 was significantly higher in DSS-treated BFΔSPT mice compared to those from BFWT mice (Figure 5G-H). These results suggest that colonic IL-18R^+^ MHC-II^+^ ILC3s may represent a specific subset of ILC3 cells that modulates IL-22 production in response to *B. fragilis*-derived sphingolipids.

### *B. fragilis-*derived sphingolipids directly inhibit ILC3-derived IL-22 in association with reduced AKT and p38 MAPK activation

To gain deeper insights into the potential interactions between bioactive sphingolipids derived from *B. fragilis* and ILC3s, we conducted *in vitro* experiments with sorted ILC3 cells. ILC3s were treated with purified *B. fragilis* sphingolipids and then stimulated with IL-23 and IL-18 to induce IL-22 production. Intriguingly, all three types of *B. fragilis* sphingolipids, including ceramide (Cers), glucosylceramide (GL-Cers), and phosphoethanolamine-ceramide (PE-Ceramide)^16^, demonstrated an inhibitory effect on the production of IL-22 by ILC3s (Figure 6A). These findings strongly suggest that *B. fragilis* sphingolipids directly inhibit IL-22 production in ILC3 cells.

**Figure 6:**
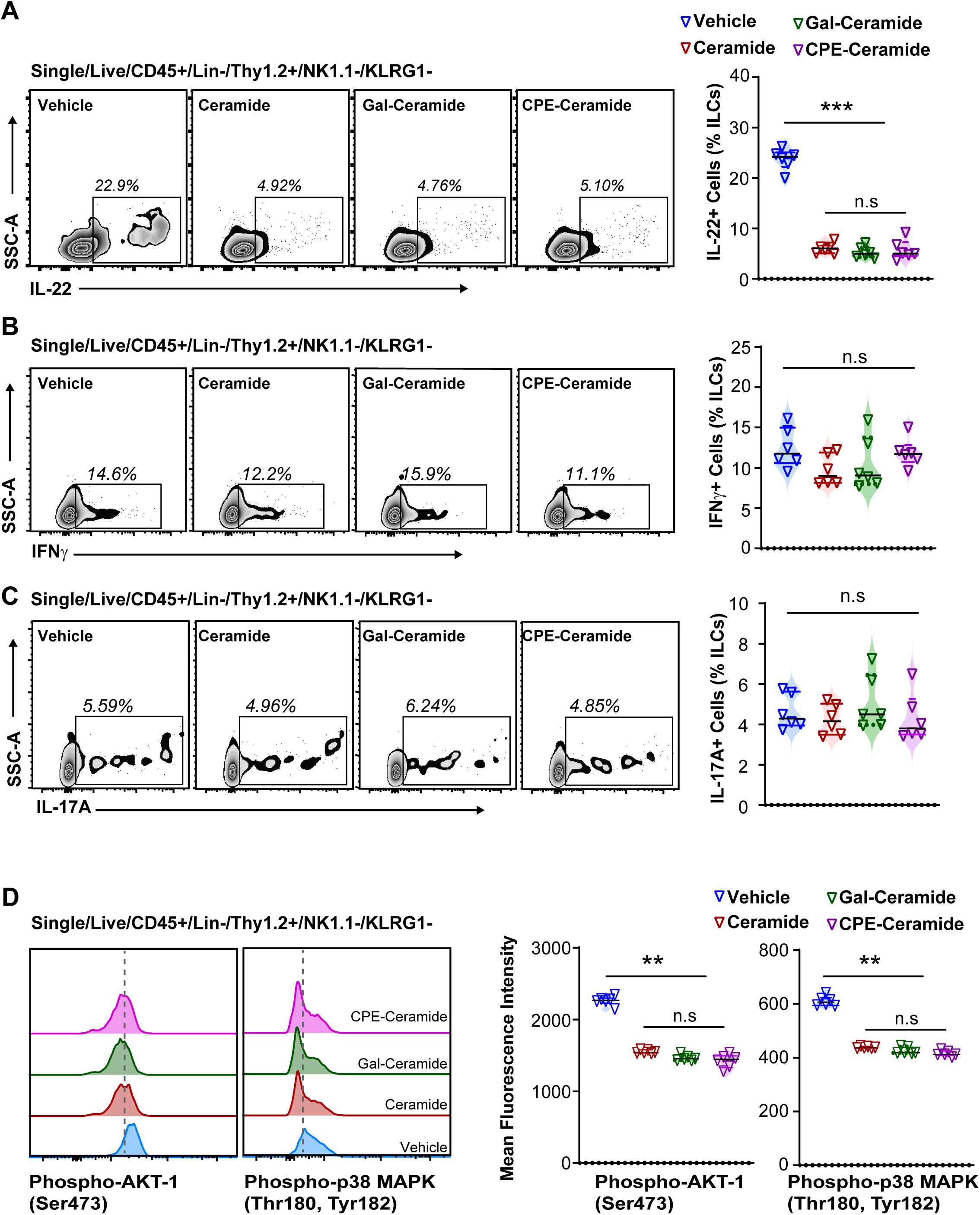
*B. fragilis*-derived sphingolipids suppress IL-22 production in ILC3 cells. (**A**) Representative flow cytometry plots and quantification of IL-22^+^ ILC3 cells after purified *B. fragilis* sphingolipids treatment and IL-23 and IL-18 stimulation. (**B**) Representative flow cytometry plots and quantification of IFNγ^+^ ILC3 cells after purified *B. fragilis* sphingolipids treatment and IL-23 and IL-18 stimulation. (**C**) Representative flow cytometry plots and quantification of IL-17A^+^ ILC3 cells after purified *B. fragilis* sphingolipids treatment and IL-23 and IL-18 stimulation (**D**) Representative flow cytometry plots of phosphorylated p38 MAPK (Thr180, Tyr182) and phosphorylated AKT-1 (Ser473) in ILC3 cells after purified *B. fragilis* sphingolipids treatment and IL-23 and IL-18 stimulation. ILC3 cells sorted from the colon and mesenteric lymph node (mLN) of germ-free mice were treated with purified sphingolipids of WT *B. fragilis* overnight. Cells were then stimulated by IL-23 and IL-18 to induce IL-22 production. ILC3 cells were sorted based on single/live/CD45^+^/Lin^-^/Thy1.2^+^/NK1.1^-^/KLRG1^-^ cells. n = 6 individual wells of 24-well-plate per group. The data represent means ± s.e.m. from two or three independent experiments. Statistical significance was determined using the nonparametric Mann-Whitney test: *p<0.05, **p<0.01, ***p<0.001.

To assess the specificity of this inhibitory effect, we examined the impact of *B. fragilis* sphingolipids on the production of other cytokines by ILC3s, including IL-17A and IFNγ. *B. fragilis* sphingolipids did not affect the production of IL-17A or IFNγ by ILC3s (Figure 6B-C). Thus, the inhibitory effect of *B. fragilis* sphingolipids appears to be specific to IL-22 production in ILC3 cells.

To elucidate the underlying molecular mechanisms by which sphingolipids modulate IL-22 production, we assessed upstream signaling pathways in ILC3 cells using phospho-flow cytometry. *B. fragilis* sphingolipids inhibited the phosphorylation of AKT-1 and p38 MAPK (Figure 6D), molecules known to regulate IL-22 production^27,41^. However, *B. fragilis* sphingolipids did not alter the phosphorylation of ERK1/2 or STAT3 signaling in ILC3s (Figure S7), pathways also known to be associated with IL-22 production in ILC3s^27,42^.

Taken together, these *in vitro* experiments suggest that *B. fragilis* sphingolipids exert their regulatory effects on IL-22 production by colonic ILC3s through targeted inhibition of AKT-1 and p38 MAPK signaling pathways.

## DISCUSSION

The gut microbiome and its metabolites play a crucial role in maintaining host mucosal homeostasis and influencing the development of IBD. However, the precise interaction between microbial metabolites and the host mucosal immune system remains largely unknown. *B. fragilis*, a prominent bacterial species in the human gut, offers an excellent model for exploring the crosstalk between the microbiome and host immune system^16,17^. Among the diverse array of microbial metabolites, sphingolipids have emerged as potential mediators that connect bioactive metabolites from the microbiome to host immune cells. Previous studies have demonstrated that *B. fragilis* sphingolipids have a beneficial effect by reducing the abundance of invariant natural killer T (iNKT) cells in murine models^16^. However, intriguingly, sphingolipids have been found to be relatively more abundant in stool samples of patients with IBD compared to that of healthy individuals^20^, suggesting a potentially detrimental role of sphingolipids in the maintenance of mucosal homeostasis. These contrasting findings highlight the complex and ambiguous role of sphingolipids in the context of the mucosal immune system. In this study, we aimed to uncover additional roles for *B. fragilis*-derived sphingolipids in host-microbiome interactions in the gut.

Our study reveals a novel and previously unrecognized role for *B. fragilis* sphingolipids in the regulation of the intestinal immune response and their contribution to the exacerbation of DSS-induced colitis (Fiugre S8). While previous studies have highlighted the beneficial effects of *B. fragilis* sphingolipids in protecting against colitis^16^, our findings suggest a more intricate role for these metabolites in the mucosal immune system. Specifically, we investigated the impact of *B. fragilis* sphingolipids on ILC3 cells, which play a crucial role in mucosal immunity responding to signals from the diet and the microbiota^22,24,25,42,43^.

We identified a novel “microbiome-ILC3 interaction” pathway, in which *B. fragilis* sphingolipids influence the ILC3-IL22 axis, leading to compromised intestinal barrier function and exacerbating DSS-induced colitis (Figure S8). We demonstrate that *B. fragilis* sphingolipids directly inhibit the production of IL-22 in colonic ILC3 cells. On the other hand, microbiota-derived short-chain fatty acids (SCFAs) enhance IL-22 production in CD4+ T cells and ILCs through GPR41-mediated HDAC inhibition, involving AhR, HIF1α, mTOR, and Stat3 pathways^44^. These findings underscore the interplay of microbial metabolites in regulating intestinal IL-22 and highlight the intricate dynamics of microbiome-host interactions in the gut.

Additionally, *B. fragilis* sphingolipids modulate IL-18 in epithelial cells, which contributes to regulating IL-22 production by ILC3 cells (Figure S8). Notably, mice mono-colonized with *B. fragilis* strains lacking sphingolipids and challenged with DSS exhibited enhanced expression of epithelial NLRC4, Caspase 1, and IL-18. Interestingly, NLRP6, which has been reported to respond to microbial metabolites and protect mice against DSS-induced colitis while promoting IL-18 production^48–50^, showed comparable expression in the epithelial cells of both WT and mutant *B. fragilis* mono-colonized mice. This suggests that the regulation of colonic IL-18 levels by *B. fragilis*-derived sphingolipids may not involve NLRP6. On the other hand, NLRP3, known for its protective role in DSS-induced colitis in myeloid cells^36^, was not detected in the colonic epithelial cells of the two groups of mono-colonized mice. These findings highlight the regulatory role of *B. fragilis* sphingolipids in modulating the NLRC4/Caspase1/L-18 axis.

Our study provides valuable insights into the specific targeting of IL18R^+^ ILC3s, particularly IL-18R^+^MHCII^+^ ILC3s, by *B. fragilis* sphingolipids to modulate IL-22 production. Additionally, we observed a reduction of colonic IL-18 levels in the presence of *B. fragilis* sphingolipids. Since colonic IL-18 is primarily produced in epithelial cells, and ILC3 MHC-II is known to interact with T cells, the modulation of IL-22 by *B. fragilis* sphingolipids may involve intricate interaction between ILC3s, intestinal epithelial cells (IEC), and lamina propria lymphocytes (LPL) (Figure S8). Further investigations are warranted to fully comprehend the complex interactions between *B. fragilis* sphingolipids and the host immune system in the context of intestinal inflammation.

ILC3 cells exert regulatory effects on various cell types within the gut mucosa through cytokines and direct cell-cell interactions, including T cells ^29,42^, iNKT cells^51^, macrophages^52^ and epithelial cells^53^. Our study demonstrates that the “microbial sphingolipids-ILC3-IL22” axis plays a pivotal role in regulating the proliferation and defense function of IECs through the STAT3 signaling pathway and downstream genes (Figure S8). Blocking IL-22 abated the protective effects fostered by the absence of *B. fragilis*-derived sphingolipid during DSS-induced colitis. These findings complement prior studies that have emphasized a critical role for IL-22 in maintaining intestinal homeostasis and facilitating tissue repair after epithelial damage^21,26,30,54–56^. Importantly, our study identifies a novel bacterial-derived modulator of IL-22.

Our study has identified a potential novel therapeutic strategy for IBD by targeting *B. fragilis* sphingolipids to enhance ILC3-derived IL-22. One goal of IBD therapy is to restore the inflamed mucosa by promoting healing and regeneration of the damaged epithelium. IL-22, derived from ILC3 cells, plays a pivotal role in mucosal healing by activating STAT3 in intestinal epithelial cells, leading to enhanced proliferation and barrier function^26,28,30,57^. Genetic studies have implicated ILC3-related genes as risk factors for IBD^58^, and decreased IL-22-producing ILC3s have been observed in the intestine of Crohn’s patients^31^. However, paradoxically, certain colonic ILC3 cells in IBD patients exhibit increased IL-22 production^59^, suggesting a complex and context-dependent role of ILC3-derived IL-22 in intestinal homeostasis and disease pathogenesis. Nonetheless, the administration of recombinant IL-22 is an intriguing therapeutic approach for IBD, and several IL-22-based clinical trials are currently underway^60,61^.

Overall, our study has uncovered a novel pathway whereby microbial sphingolipids modulate both ILC3s and epithelial cells to regulate intestinal inflammation (Figure S8). Further exploration and validation of the “bacterial sphingolipids-ILC3-IL22” axis in human IBD may offer a novel therapeutic approach for managing this complex and challenging disease.

## Methods

### Mice

All animal experiments and procedures adhered to the National Research Council’s Guide to the Care and Use of Laboratory Animals and were conducted in compliance with Institutional Animal Care and Use Committee-approved protocols 14-11-2802, 17-11-3556R, and 20-11-4280R (IACUC, Boston Children’s Hospital). C57BL/6 background wild-type mice were obtained from the Jackson Laboratories and maintained under specific pathogen-free (SPF) or germ-free (GF) conditions. GF and monocolonized mice were bred and maintained in vinyl isolators in the Animal Resources at Children’s Hospital (ARCH). Monocolonized mice (BFWT and BFΔSPT mice) were generated, and their offspring were maintained as previously described^16^. Stool samples from GF and monocolonized mice were routinely collected and tested for sterility and exclusion of other bacterial contamination using aerobic and anaerobic conditions culture and 16s PCR accompanied by sequencing.

DSS-induced colitis was induced in the isolators by administering 1.5% colitis-grade DSS (36,000 – 50,000 Da, MP Bioscience) in water, which was prepared and filtered through a 0.22 μm filter system in a sterile laboratory hood. The DSS water and all other necessary supplies were sterilized before being transferred into the isolator. At indicated time points, the mice were weighed and evaluated for diarrhea and rectal bleeding consistently at the same time of day.

### Organoids

Organoids were derived from isolated colonic crypts of germ-free C57/BL6 mice using previously established methods^62^. Roughly 500 crypts were plated in 24-well plates within a 50 μl drop of Matrigel, overlaid with 500 μl of WENR medium. This medium consisted of basal crypt media (Advanced DMEM/F12, penicillin/streptomycin, 10 mM HEPES, 2 mM Glutamine) supplemented with 1× B27 (Gibco, 17504-044), 1× N2 (Gibco, 17502-048), 50 ng/ml rmEGF (Peprotech, 315-09), 50% L-WRN-CM (conditioned medium) (v/v), 10 μM Rock inhibitor Y-27632 (Sigma, Y0503), and 3 μM CHIR99021 (Sigma, SML1046), with a final FBS concentration of 10%. Media was refreshed every 2 days, and organoids were passaged every 4 days, with three passages conducted before the start of experiments. Organoids were treated with a complete medium containing either total *B. fragilis* lipids or purified sphingolipids and incubated at 37°C with 5% CO2 in a humidified incubator to support organoid formation and growth. Caution was exercised to minimize disturbance during the initial phase of organoid development. Subsequent to 72-96 hours of treatment, images were captured using the Bio Tek Cytation 5 automated microscope, and cells were harvested from the recovered organoids for flow cytometry or protein extraction analysis.

### Bacterial culture and Lipid extraction

*Bacteroides. fragilis* (WT, NCTC9343; Mutant, BF9343-Δ3671) was cultured according to previously established methods^16^. Briefly, frozen stocks were streaked onto a Blood Agar plate and incubated for 24 hours at 37 °C in an anaerobic system. A single colony was then selected and grown in 5 mL of the rich medium within an anaerobic chamber. To collect samples for lipid extraction, *B. fragilis* was cultivated in 1 liter of medium to an OD of 1.0, centrifuged, washed twice with phosphate-buffered saline (PBS), and stored at -80 °C until extraction.

Lipids from *B. fragilis* were extracted using a modified version of a previously established method^16^. Bacterial sphingolipids were isolated from *B. fragilis* anaerobic cultures (either BFWT or BFΔSPT) using a base-treated lipophilic extraction technique. After stirring the washed bacterial pellets in chloroform/methanol (2:1) for five hours, the mixture was concentrated to dryness using a rotovap. Subsequently, the material was treated with 0.01 N NaOH in methanol for five hours, followed by dilution with methanol, chloroform, and 1 N NaCl (aq) to achieve a final ratio of 4:2:1.6. After stirring for four hours, additional chloroform was added to the mixture in a separatory funnel. The bottom (chloroform) phase was collected and separated through multiple iterations, repeating the separation process with chloroform. Further purification was accomplished using a flash silica gel chromatography method, employing eluents ranging from 8:1 to 1:2 chloroform/methanol, and thin-layer chromatography. The purified lipids were dissolved in DMSO at a concentration of 250 μg/mL. Prior to treatment, the lipids were diluted in a cell culture medium to a final concentration of 250 ng/mL.

### Isolation of intestinal epithelial cells (IEC), intraepithelial lymphocytes (IEL), and lamina propria lymphocytes (LPL)

Colons were dissected, and the remaining fat was removed before cleaning the contents with cold PBS. Intestines were inverted to expose the epithelial layer, which was then allowed to interact with the buffer. Mucus was removed by shaking the intestines in ice-cold PBS. The dissociation of the epithelial layer, including epithelial cells and IEL, was achieved by incubating on a shaker in extraction buffer (PBS with 2% FBS, 2 mM EDTA, 1 mM DTT) for 25 minutes at 37 °C. The remaining samples were washed with PBS and used for LPL digestion.

To extract IEC and IEL, the epithelial fraction was passed through a 40 μm cell strainer (Falcon), centrifuged, and washed twice with PBS containing 2% FBS. Leukocytes and epithelial cells were separated and enriched by 40%/80% Percoll (Cytiva) gradient centrifugation.

To isolate LPL, the intestine was cut into 1 mm sections and digested with digestion buffer (RPMI with 5% FBS, 1.5 mg/ml Collagenase II, 0.5 mg/ml Dispase) on a shaker for 45 minutes at 37°C. The resulting fraction was then passed through a 40 μm cell strainer (Falcon), centrifuged, and washed twice with RPMI with 5% FBS. Leukocytes were further enriched by 40%/80% Percoll (Cytiva) gradient centrifugation. Isolated single-cell suspensions were used for flow cytometry analysis, cell sorting, RNA extraction, and protein extraction.

### Colon explant culture and ELISA

The colon was dissected, and its contents were removed using ice-cold PBS with Penicillin-Streptomycin (100 U/ml). Then, a 1 cm section of the distal colon was cut and opened longitudinally. It was then incubated for 1 hour at 37 °C in RPMI 1640 supplemented with FBS (10%) and Penicillin-Streptomycin (100 U/ml). The colon explant was then transferred to a well of a 24-well plate with fresh RPMI 1640 supplemented with FBS (10%) and Penicillin-Streptomycin (100 U/ml) and incubated for 24 hours at 37 °C. The medium was collected and stored at -80 °C for cytokine analysis.

Cytokines, including TNFa, IFN-γ, MCP-1, IL-1β, IL-10, IL-6, IL-22, IL-23, and IL-17A, were measured using the Biolegend ELISA MAX kit following the manufacturer’s instructions. IL-18 was assayed using the IL-18 Mouse ELISA Kit (Invitrogen) according to the instructions.

### Flow cytometry

Single-cell suspensions were loaded onto a 96-well plate and washed with in ice-cold PBS. To exclude dead cells, staining with eBioscience™ Fixable Viability Dye eFluor™ 506 were performed. Fc receptors were blocked using TruStain FcX™ PLUS (anti-mouse CD16/32) Antibody from Biolegend. Surface staining was carried out by incubating cells with staining antibodies in FACS buffer (PBS with 2% FBS and 1 mM EDTA) on ice. After washing with FACS buffer, cells were fixed with Cytofix/Cytoperm™ from BD Bioscience on ice and permeabilized with Foxp3/Transcription Factor Staining Buffer Set from eBioscience, Invitrogen. Intracellular staining was performed using staining antibodies in the permeabilization buffer, followed by washing with the same buffer. Finally, cells were refixed with Cytofix/Cytoperm™.

For cytokine staining, cells were incubated for 6 hours in RPMI 1640 with 10% FBS, PMA (phorbol 12-myristate-13-acetate) at 50 ng/mL, ionomycin at 750 ng/mL, and protein transport inhibitor (Brefeldin A) at 10 μg/mL. Most of the staining antibodies were purchased from Biolegend, BD Bioscience, and Thermo Fisher Scientific. Various antibodies were used for staining, including CD45 (30-F11), CD90.2 (30-H12), Lineage cocktail (145-2C11, RB6-8C5, RA3-6B2, Ter-119, M1/70), KLRG1 (2F1/KLRG1), CD3 (17A2), CD4 (GK1.5), CD8a (53-6.7), CD19 (6D5), CD127 (A7R34), CCR6 (29-2L17), NKp46 (29A1.4), IL-22 (Poly5164), NK1.1 (PK136), RORγt (B2D), Phospho-ERK1/2 (Thr202, Tyr204) (MILAN8R), Phospho-AKT1 (Ser473) (SDRNR), Phospho-STAT3 (Tyr705) (4NIT4KK), Phospho-p38 MAPK (Thr180, Tyr182) (MILAN8R), B220 (RA3-6B2), T-bet (4B10), IL-17A (TC11-18H10.1), IFN-γ (XMG1.2), TCR γδ (UC7-13D5), FOXP3 (FJK-16s), GATA3 (TWAJ), Ep-CAM (G8.8), and Ki67 (11F6). Surface antibodies were diluted 1:200, intracellular antibodies were diluted 1:100, and phospho antibodies were diluted 1:20.

All flow cytometry experiments were conducted on the LSRFortessa™ Cell Analyzer from the Harvard Digestive Diseases Center, using the FACS Diva software from BD Bioscience. The FACS data were analyzed with FlowJo V10 software from Tree Star.

### ILC3 cell sorting and stimulation

ILC3 cells (CD45+Lin-CD90.2+CD127+KLRG1-) were sorted using the FACSMelody Cell Sorter and the FACSChorus™ software (BD Bioscience). The cells were then incubated in DMEM (high glucose) with FBS (10%), sodium pyruvate (1 mM), HEPES (10 mM), MEM nonessential amino acids, 2-Mercaptoethanol (BME) (55 μM), and Penicillin-Streptomycin (100 U/ml) with IL-23 (20 ng/ml, Biolegend), in the presence of purified *B. fragilis* Sphingolipids (Cers, GL-Cers, PE-Cers; 250 ng/ml) or DMSO for 4 hours at 37°C. To analyze IL-22, a protein transport inhibitor (Brefeldin A) (10 μg/ml) was added.

### RNA extraction, qPCR, and RNA sequencing

The RNA extraction process involved lysing mouse colonic IEC or cultured cells using RLT buffer (QIAGEN) and extracting total RNA using the RNeasy Mini Kit (QIAGEN) following the manufacturer’s instructions. For qPCR, cDNA was synthesized using the QuantiTect Reverse Transcription Kit (QIAGEN), and real-time PCR was performed on a LifeTechnologies QuantStudio 7 real-time PCR system using SYBR™ Green PCR Master Mix (Applied Biosystems). Gene expression levels were normalized to Actb or Gapdh, and data analysis was performed using the ΔΔCt method. For RNA sequencing, libraries were prepared using the NEBNext® Ultra™ II Directional RNA Library Prep with Sample Purification Beads (New England Biolabs) according to the provided instructions. The quality of RNA and cDNA libraries was assessed using a TapeStation. Sequencing was conducted on the Illumina NextSeq 500 platform at the Biopolymers Facility at Harvard Medical School. After quality control checks, the raw RNA-seq data was processed, including read alignment, quantification of gene expression, and differential expression analysis, which were performed using RStudio. The results were visualized using the EnhancedVolcan R package.

### Western Blot

Mouse colonic IECs or cultured cells were lysed using RIPA buffer (Cell Signaling) to extract proteins. Protein samples were separated using 4%-15% Mini-Protean TGX Precast Protein Gels. For detection, the following primary antibodies were used: anti-phospho-Stat3 (Tyr705) (D3A7, Cell Signaling), anti-Stat3 (124H6, Cell Signaling), and anti-β-Actin (AC-15, Sigma-Aldrich). Horseradish peroxidase-conjugated anti-mouse or anti-rabbit secondary antibodies were used for detection.

### Statistical Analysis

Statistical analyses were conducted using GraphPad Prism V9 software. A two-tailed, nonpaired Student’s t-test or nonparametric Mann-Whitney test was used as specified in each figure to determine the P-value. In the figures, statistical significance is denoted by asterisks as follows: *P < 0.05, **P < 0.01, ***P < 0.001.

## Supporting information

supplementary Table-1--RNA seq results-Differentially expressed genes in colonic epithelial cells from DSS-treated BFWT and BFdeltaSPT mice

supplementary Table-2--RNA seq results-Up-regulated genes in colonic epithelial cells from DSS-treated BFdeltaSPT mice

supplementary Table-3--RNA seq results-Down-regulated genes in colonic epithelial cells from DSS-treated BFdeltaSPT mice

## Acknowledgments

We extend our gratitude to Ellen Deniken from Genentech, Inc (South San Francisco) for generously providing the IL-22 antibody (IL-22:9592). We also acknowledge the valuable assistance of Andrew Kwong and Faith Taliaferro from Dr. Jose Ordovas-Montanes’s lab for their contributions to RNAseq data analysis. This work was supported by P30DK03485 (S.B.S; B.B), the Wolpow Family Chair in IBD Treatment and Research (S.B.S), the Rainin Foundation (S.B.S) and the National Natural Science Foundation of China (31401204, U1532269 to B.B.).

## Authors contributions

Planning and conceptualization, B.B., D.A., D.L.K., S.B.S.; Experiments and Procedures, B.B., Y.W., X.S., M.W., I.P., Y.T., A.R., J.B., and X.C.; Sphingolipids Purification, P.B., and E.B.; Provision of Key Resources, D.L.K., W.L., M.W., J.L., J.T., C.W., and S.B.S.; Writing and Editing Manuscript, B.B., J.O., B.H., and S.B.S. with review from all authors.

## Figure legends for supplementary figures

**Figure S1:**
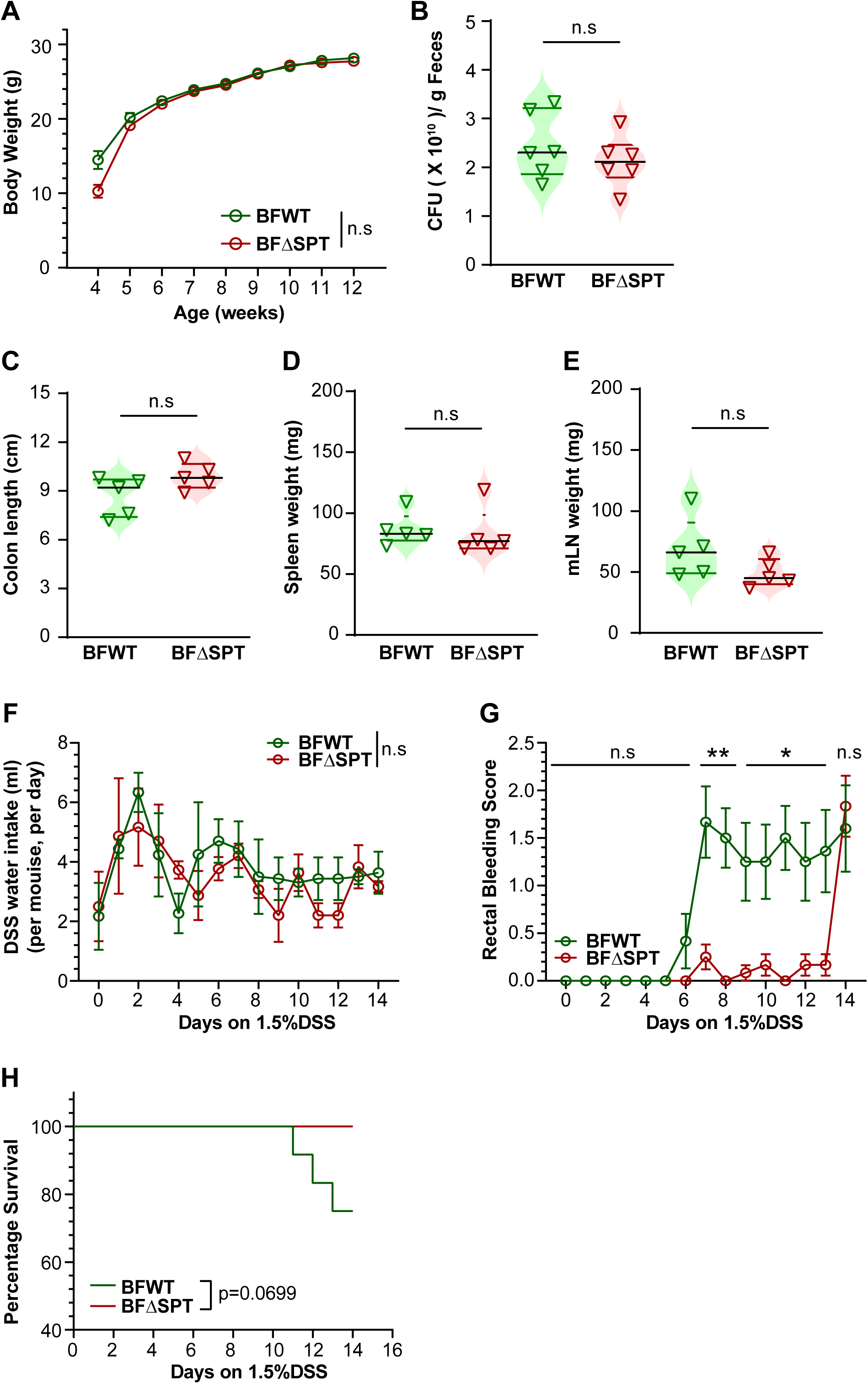
Effects of *B. fragilis*-derived sphingolipids at baseline and following DSS-induced colitis. (**A**) Body weight after weaning. (**B**) Colony forming units (CFU) of *B. fragilis* in fecal samples from mono-colonized mice. (**C**) Colon length, (**D**) spleen weight, and (**E**) mesenteric lymph node (mLN) weight of 12-week-old mono-colonized mice. Mono-colonized mice were treated with 1.5% DSS in their drinking water for 14 days, and the following parameters were monitored: (**F**) DSS water intake (**G**) rectal bleeding scores, and (**H**) survival rate. The experiment was conducted with 5-7 mice per group, and the data represent two or three independent experiments, presented as means ± standard error of the mean (s.e.m). The data represent two or three independent experiments, shown as means ± s.e.m., with n = 5-6 mice per group. Statistical significance was determined using the nonparametric Mann-Whitney test, where *p<0.05, **p<0.01, and ***p<0.001.

**Figure S2:**
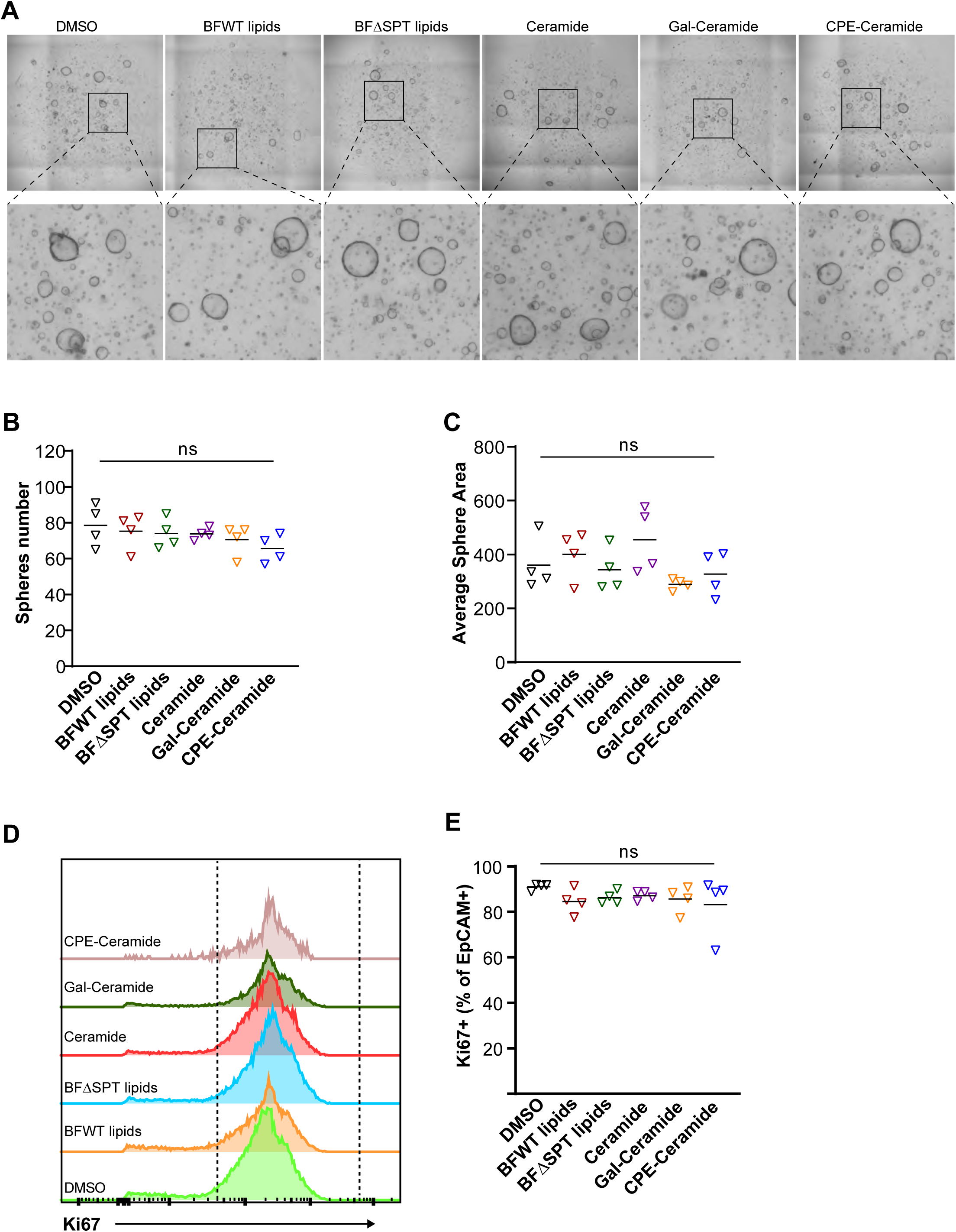
Effect of *B. fragilis*-derived sphingolipids on organoid growth and cell proliferation. (**A**) Representative images of organoids and (**B**) quantification of the number of spheres formed, and (**C**) average sphere area of organoids treated with total lipids or purified sphingolipids derived from *B. fragilis*. (**D**) Histogram of fluorescence intensity and (**E**) quantification of the percentage of Ki67^+^ cells in cells extracted from organoids treated with total lipids or purified sphingolipids from *B. fragilis*. Data are presented as means ± standard error of the mean (s.e.m.) from two or three independent experiments. Statistical significance was determined using the nonparametric Mann-Whitney test, where *p<0.05, **p<0.01, and ***p<0.001.

**Figure S3.**
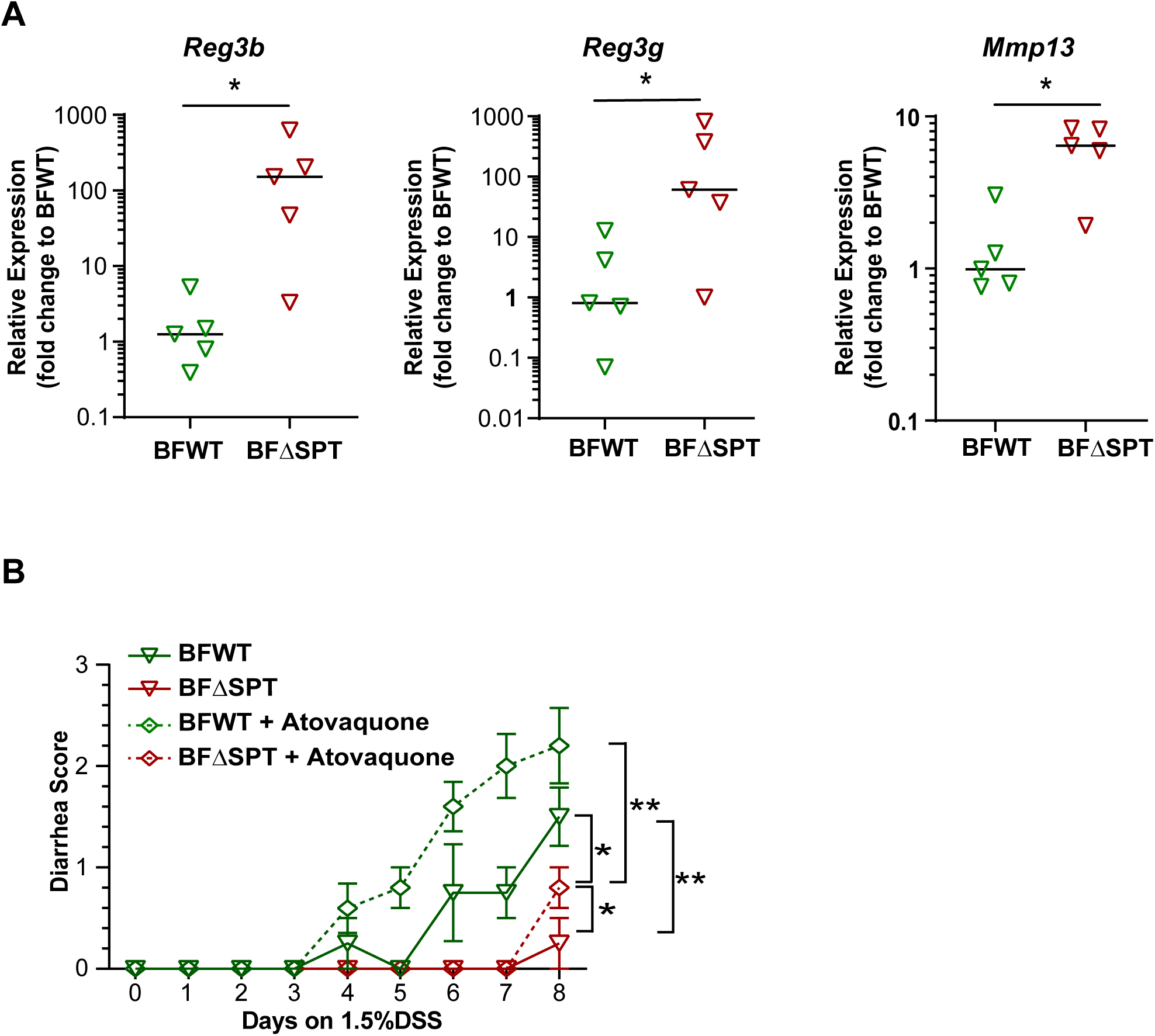
Quantification of STAT3 target genes and effect of atovaquone treatment on diarrhea scores in mono-colonized mice during DSS treatment. (**A**) Quantitative Reverse Transcription PCR (qRT-PCR) analysis of Reg3b, Reg3g, and Mmp13 expression in colonic epithelial cells from mono-colonized mice treated with 1.5% DSS for 14 days. (n = 5 mice per group.) (**B**) Diarrhea scores were examined during the DSS treatment. Mono-colonized mice received either atovaquone or PBS gavage and were challenged with DSS in their drinking water for 8 days. The experiment was conducted with 5-7 mice per group, and the data represent two or three independent experiments, presented as means ± standard error of the mean (s.e.m.). Statistical significance was determined using the nonparametric Mann-Whitney test, with *p<0.05, **p<0.01, and ***p<0.001 indicating significant differences.

**Figure S4.**
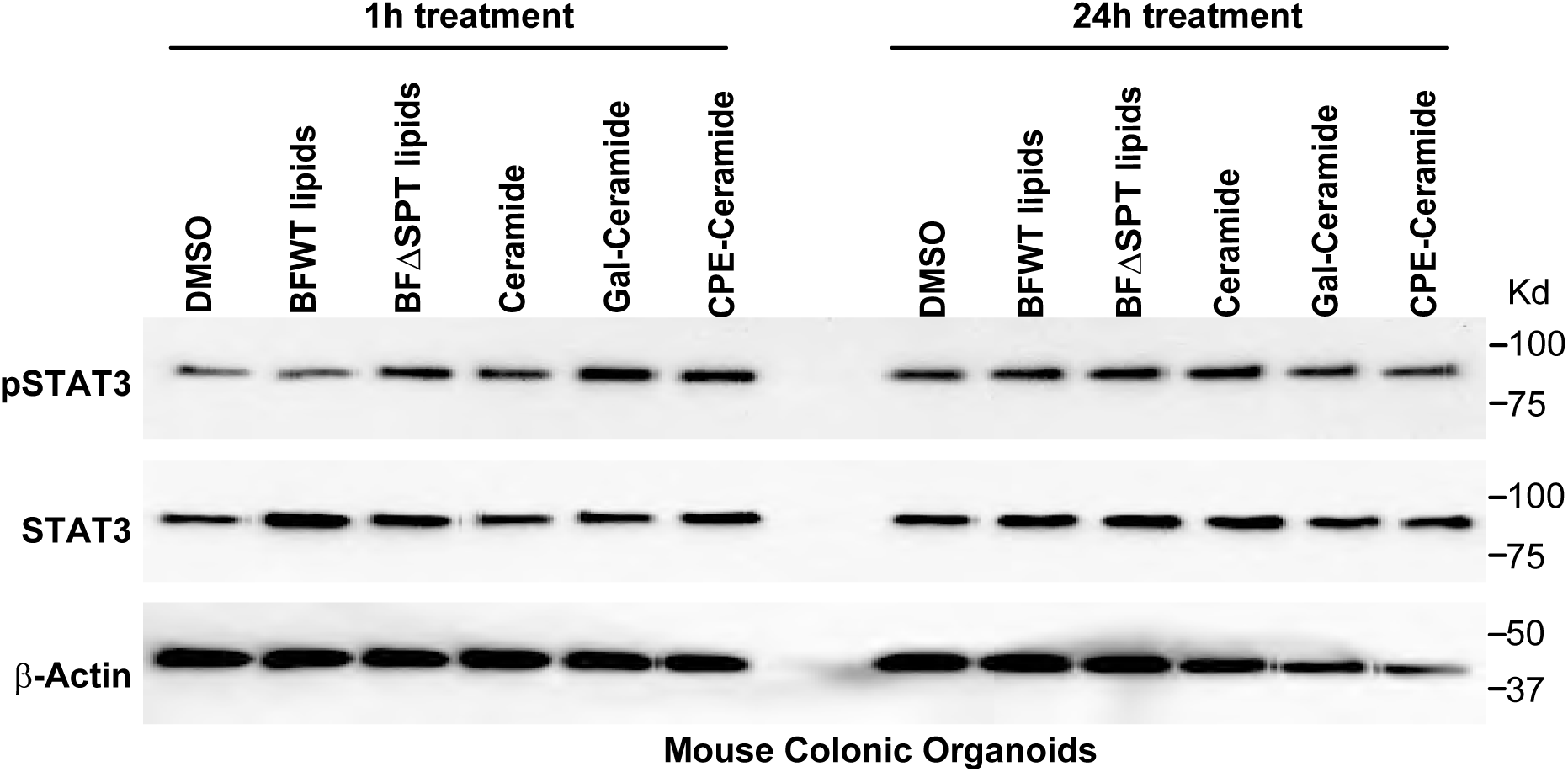
*B. fragilis*-derived sphingolipids do not directly modulate epithelial cell STAT3 activation. Representative Immunoblot phosphorylated and total STAT3 in mouse colonic organoids treated with total lipids extracted from WT or mutant *B. fragilis* or purified *B. fragilis* sphingolipids. Data represent two or three independent experiments shown as the means± s.e.m. The nonparametric Mann-Whitney test was used to determine statistical significance. *p<0.05, **p<0.01, ***p<0.001.

**Figure S5.**
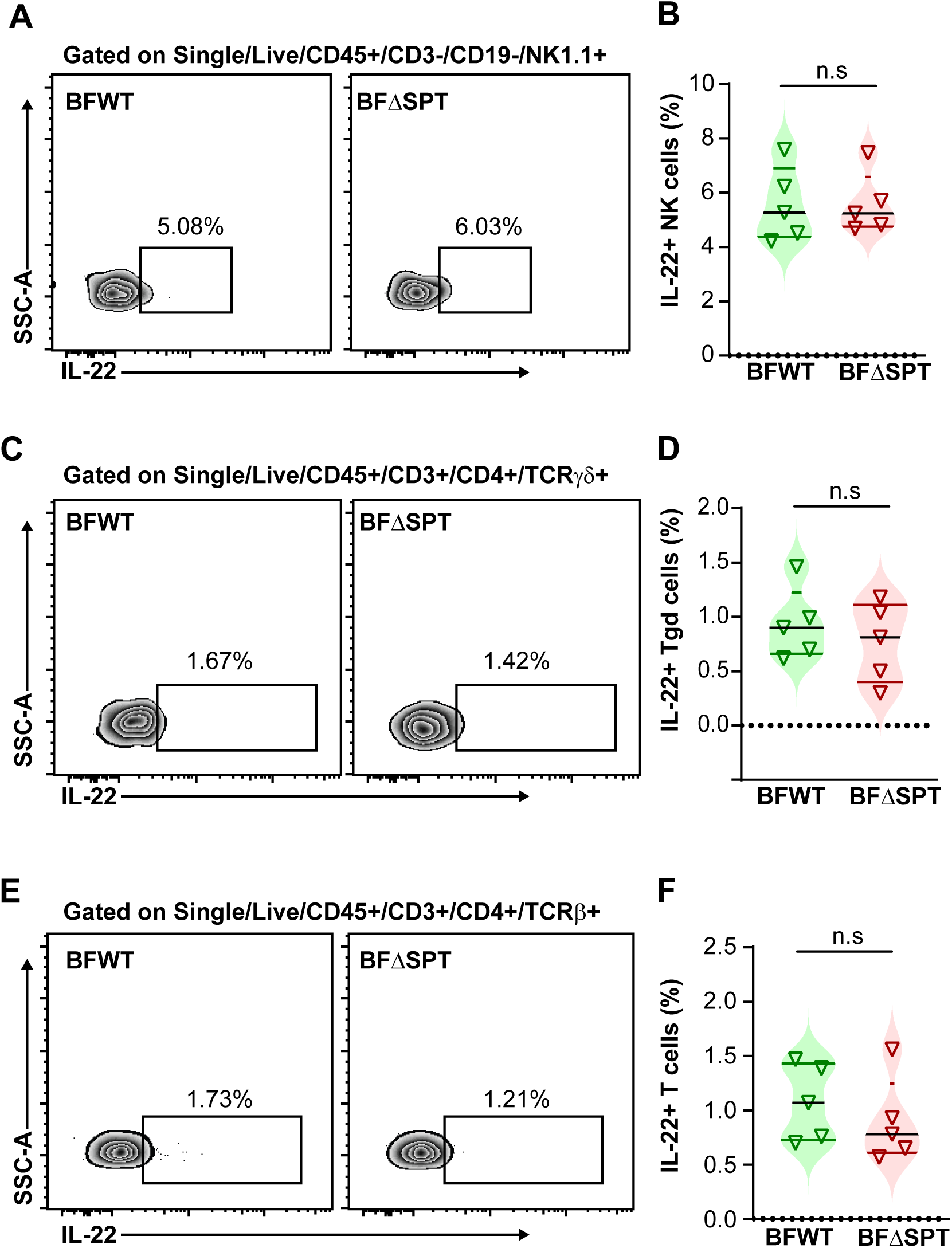
*B. fragilis*-derived sphingolipids do not affect IL-22 production in NK and T cells. (**A**) Representative flow cytometry plots and (**B**) quantification of IL-22^+^ cells in NK cells in the colon of mono-colonized mice after 7-day DSS treatment. (**C**) Representative flow cytometry plots and (**D**) quantification of IL-22^+^ cells in γδT cells in the colon of mono-colonized mice after 7-day DSS treatment. (**E**) Representative flow cytometry plots and (**F**) quantification of IL-22^+^ cells in TCRβ+ T cells in the colon of mono-colonized mice after 7-day DSS treatment. Mono-colonized mice were challenged with 1.5% DSS in drinking water for 7 days. n = 5 mice per group. Data represent two or three independent experiments shown as means ± s.e.m.. Statistical significance was determined by the nonparametric Mann-Whitney test. *p<0.05, **p<0.01, ***p<0.001.

**Figure S6.**
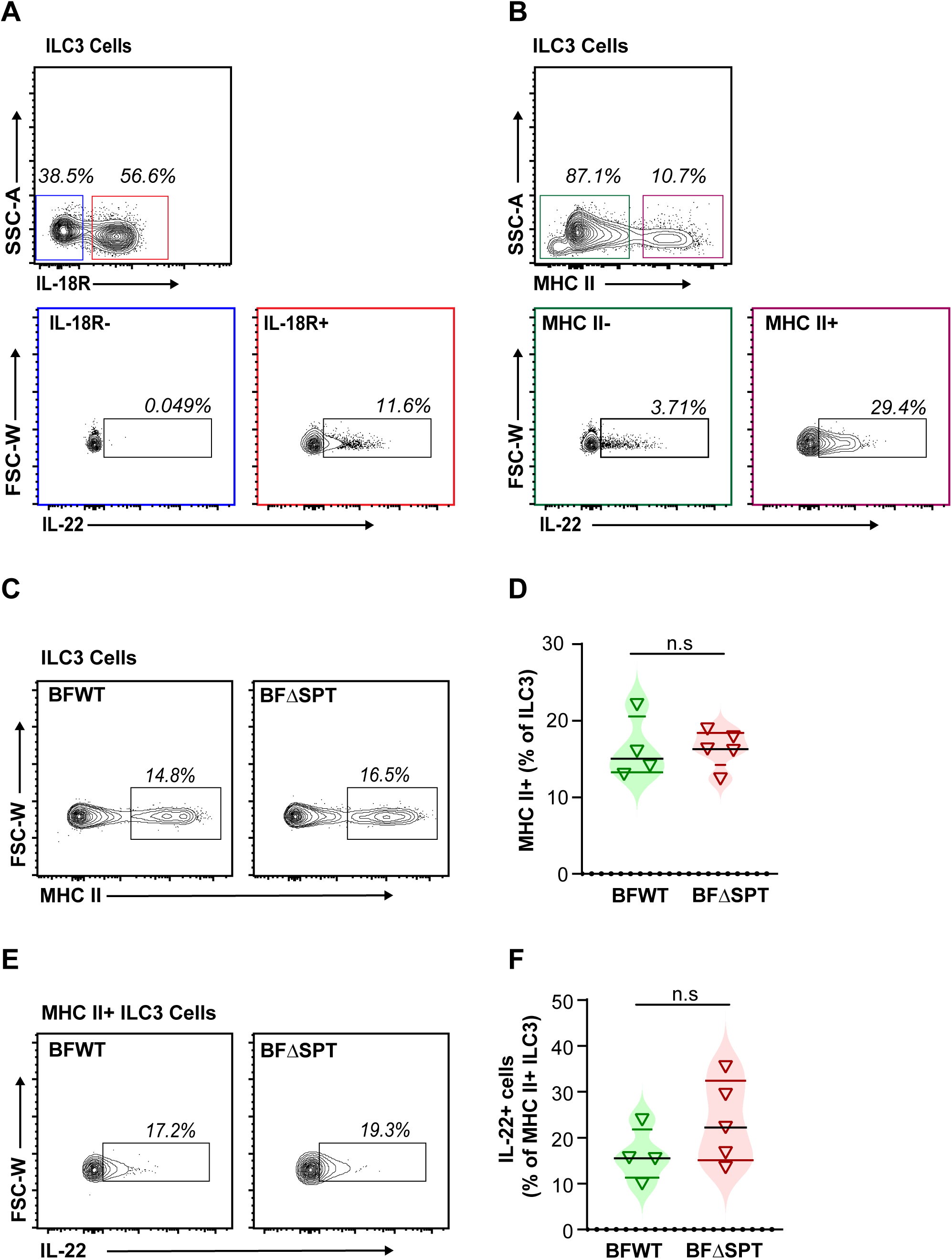
*: B. fragilis*-derived sphingolipids modulate ILC3-derived IL-22 by targeting IL18R+ MHCII+ ILC3 cells. (**A**) Representative flow cytometry plots of IL-18R^+^ / IL-18R^-^ ILC3 cells, as well as IL-22^+^ cells in IL-18R^+^ / IL-18R^-^ ILC3 cells from the colon of mono-colonized mice after 7 days of 1.5% DSS challenge. (**B**) Representative flow cytometry plots of MHC II ^+^ / MHC II ^-^ ILC3 cells, as well as IL-22^+^ cells in MHC II ^+^ / MHC II ^-^ ILC3 cells from the colon of mono-colonized mice after 7 days of 1.5% DSS challenge. (**C**) Representative flow cytometry plots and (**D**) quantification of MHCII^+^ ILC3 cells in the colon of mono-colonized mice after 7 days of 1.5% DSS challenge. (**E**) Representative flow cytometry plots and (**F**) quantification of IL-22^+^ cells in MHCII^+^ ILC3 cells in the colon of mono-colonized mice after 7 days of 1.5% DSS challenge. ILC3 cells are gated on single/live/CD45^+^/Lin^-^/Thy1.2^+^/IL7R^+^/RORγt^+^ cells. A-C, n = 8 individual mice per group; D-M, n = 4-5 individual mice per group. Data represents means± s.e.m. from two or three independent experiments. The nonparametric Mann-Whitney test determined statistical significance (*p<0.05, **p<0.01, ***p<0.001).

**Figure S7:**
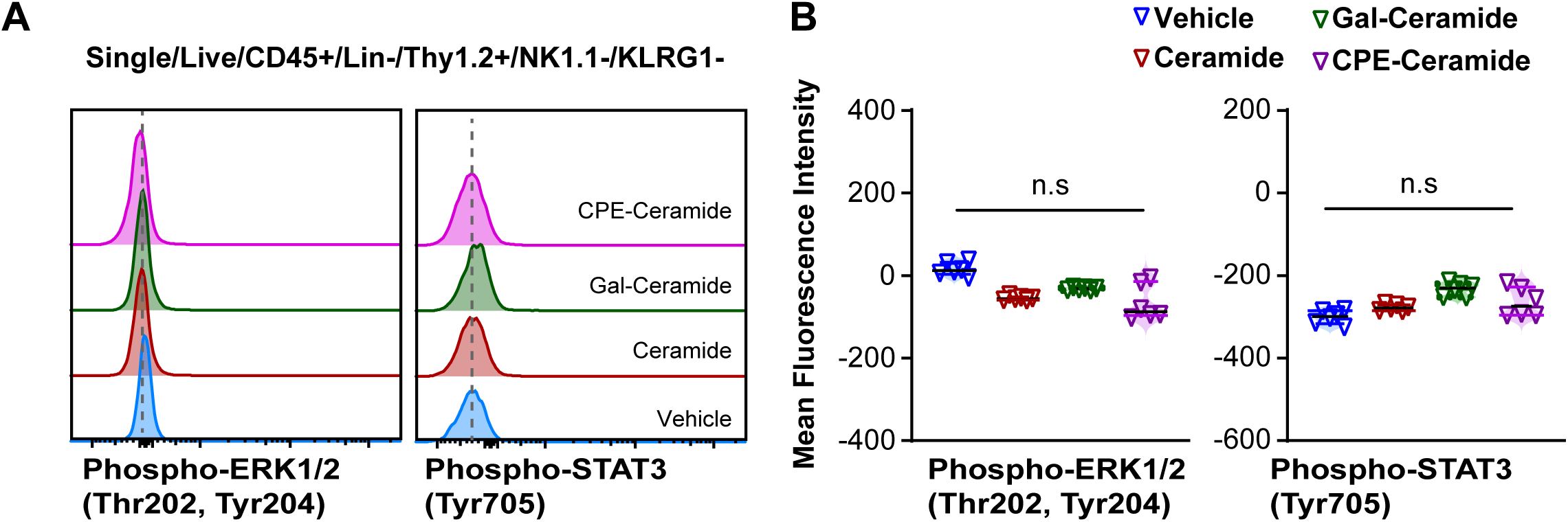
*B. fragilis*-derived sphingolipids suppress IL-22 production in ILC3 cells. (**A**) Representative flow cytometry plots and (**B**) quantification of phosphorylated ERK1/2 (Thr202, Tyr204) and phosphorylated STAT3 (Tyr705) in ILC3 cells after purified *B. fragilis* sphingolipids treatment and IL-23 and IL-18 stimulation. ILC3 cells were sorted based on single/live/CD45^+^/Lin^-^/Thy1.2^+^/NK1.1^-^/KLRG1^-^ cells from the colon and mesenteric lymph node (mLN) of germ-free mice. Following sorting, the cells were treated with purified sphingolipids from WT *B. fragilis* overnight. Subsequently, cells were then stimulated with IL-23 and IL-18 to induce IL-22 production. ILC3 cells were sorted n = 6 individual wells of 24-well-plate per group. The data represent means ± s.e.m. from two or three independent experiments. Statistical significance was determined using the nonparametric Mann-Whitney test: *p<0.05, **p<0.01, ***p<0.001.

**Figure S8:**
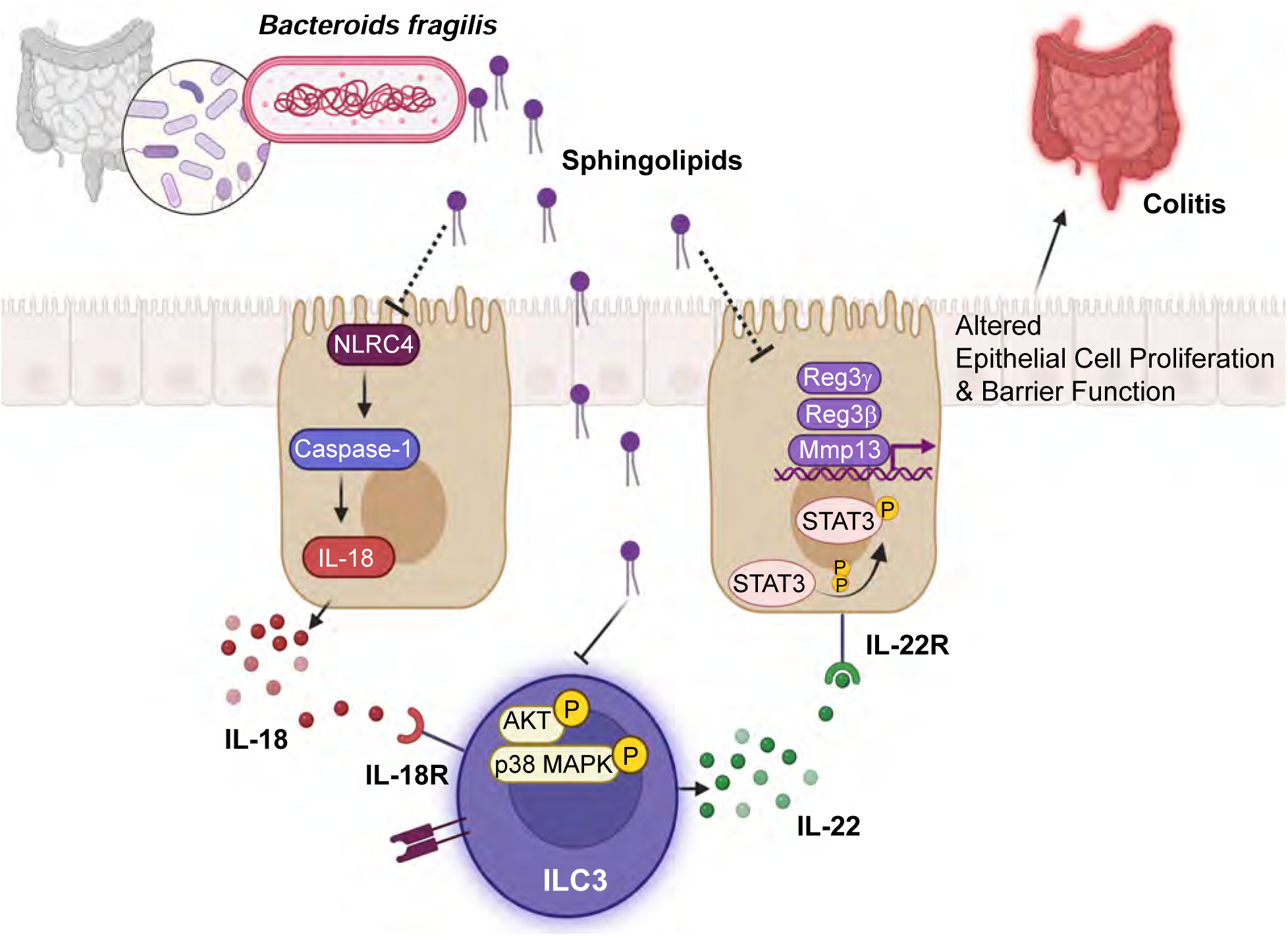
Model illustrating the interaction between *B. fragilis*-derived sphingolipids and the host mucosal immune system. *B. fragilis* sphingolipids modulate the immune response by restraining NLRC4 and Caspase-1 activation, leading to reduced IL-18 levels in the gut. Simultaneously, *B. fragilis* sphingolipids interact with a specific subset of immune cells known as IL-18R+ MHCII+ ILC3 cells. This interaction results in the suppression of AKT and p38 MAPK activity in these ILC3 cells, leading to a reduction in IL-22 production. The decrease in IL-22 levels in the gut results in compromised activation of STAT3 in the intestinal epithelial cells. Consequently, epithelial barrier function and wound healing are compromised, contributing to the development of colitis. Overall, the model demonstrates the intricate interplay between *B. fragilis*-derived sphingolipids, the gut immune system, and epithelial cells, highlighting the critical role of these sphingolipids in modulating mucosal homeostasis and influencing colitis development.

## Notes

### Competing Interest Statement

The authors have declared no competing interest.

